# Can carcasses cause environmental nucleic acid pollution? Implications for interpreting environmental RNA/DNA survey results

**DOI:** 10.64898/2026.10.08.757455

**Authors:** Yasuaki Inoue, Kaede Miyata, Taijun Myosho, Masayuki Yamane, Hiroshi Honda

## Abstract

Environmental nucleic acid (eRNA and eDNA) metabarcoding is a promising tool for biodiversity assessment as it supports low-cost, noninvasive biological surveys. A critical challenge for eRNA/eDNA analyses is accounting for nucleic acid contamination that can compromise the accuracy of biomonitoring results. The primary aim of the current study was to examine carcasses as potential sources of eRNA/eDNA contamination, a contribution that remains poorly understood. Using Japanese medaka (*Oryzias latipes*), we quantified the detectability and persistence of carcass-derived eRNA/eDNA in water through two complementary experiments: (i) a shedding study comparing eRNA/eDNA release from live and dead individuals and (ii) a decay experiment measuring temperature-dependent degradation from 4 to 34 °C. Dead fish released detectable amounts of both eDNA and eRNA, indicating that eRNA-based surveys cannot fully eliminate carcass-derived signals. Arrhenius-based analysis showed that eRNA degradation was more temperature-dependent than eDNA degradation. These results suggest that eRNA may improve temporal resolution for recent organismal presence under warm conditions, whereas carcass-derived eRNA may persist under cold conditions and complicate interpretation of positive detections. Overall, eRNA may help reduce carcass-related false positives, particularly in warm conditions, but should be paired with temperature-aware sampling strategies and temporal replication during colder periods.

## INTRODUCTION

As global biodiversity loss accelerates, sensitive and cost-effective monitoring methods are increasingly needed, making environmental nucleic acid (eNA) approaches particularly valuable. Environmental DNA (eDNA) is now widely used in aquatic biomonitoring as it can detect target taxa more sensitively than conventional capture- or observation-based surveys, thereby supporting biodiversity assessment and environmental management decisions.^1–4^ However, eDNA detected in water does not necessarily indicate the current presence of living organisms, as nucleic acids can also originate from nonliving sources such as carcasses and anthropogenic effluents, including wastewater-treatment-plant discharges and fish-market wastewater. ^5–7^

Environmental RNA (eRNA) has also gained attention as a biomonitoring tool because its rapid degradation may more closely reflect recent organismal presence.^7–13^ We previously reported that domestic wastewater and wastewater-treatment-plant effluent can generate false-positive signals in eNA-based analyses, whereas eRNA-based analyses may help reduce these signals.^7^ However, whether eRNA-based analyses can similarly reduce carcass-derived false positives remains unclear.

Two issues are central to evaluating carcass effects on eRNA-based monitoring. First, it is unclear whether carcasses release eRNA into aquatic environments. Carcasses can release more eDNA than living fish, suggesting that carcass-derived signals cannot be fully excluded from eDNA analyses. ^14^ If decomposing organisms release substantial nucleic acids, these signals may confound the interpretation of positive detections.^15–17^ Second, if carcasses release eRNA, its decay dynamics in water must be characterized. Rapid degradation would limit the influence of carcass-derived eRNA, whereas prolonged persistence could increase false-positive risk. Although eRNA is often assumed to decay faster than eDNA, reported decay rates vary among studies.^18–21^ Although two studies have compared eRNA and eDNA decay across temperatures, their temperature ranges were relatively narrow (10–30 °C). Hence, decay at low, winter-relevant temperatures remains poorly understood.^20,21^ Clarifying which molecule decays faster, and under which temperature conditions, could inform waterbody- and season-specific monitoring strategies.

Accordingly, in the current study, we quantified the detectability and persistence of carcass-derived eRNA using Japanese medaka (*Oryzias latipes*) in two complementary experiments: a release experiment comparing live and dead fish and a decay experiment conducted at 4, 14, 24, and 34 °C. We also applied Arrhenius-based kinetic analysis to quantify the temperature dependence of eRNA and eDNA decay and compare their mechanistic differences. ^22–26^ We tested two hypotheses: (i) dead *O. latipes* do not release detectable eRNA and (ii) eRNA decays faster than eDNA across temperatures. Together, these experiments allowed us to evaluate whether eRNA can reduce carcass-derived signals and clarify the utility and limitations of eRNA monitoring.

## METHODS

### eRNA/eDNA shedding from *Oryzias latipes*

*O. latipes* used in this study was obtained from the National Institute for Environmental Studies (Japan) and maintained in our laboratory for more than three generations. Fish were reared under constant conditions (24 ± 1 °C, 16 h light:8 h dark photoperiod) and fed brine shrimp twice daily. To minimize fecal effects, fish were not fed from 24 h before the experiment until its completion. Before the experiments, glass aquaria (20 × 15 × 20 cm; length, width, and height) were washed with detergent, 1% sodium hypochlorite solution, and ethanol, then rinsed several times with distilled water. Each aquarium was filled with 3 L of dechlorinated water, and water temperature was maintained at 24 ± 1 °C using an open-circuit chiller (Taitec Corp., Saitama, Japan). Fifteen live fish were added to each aquarium and acclimated under continuous aeration (Figure 1a). Time 0 (*t* = 0) was defined as immediately after fish were introduced. We prepared triplicate aquaria for each sampling time point, along with a negative control aquarium containing dechlorinated water without fish.

**Figure 1.**
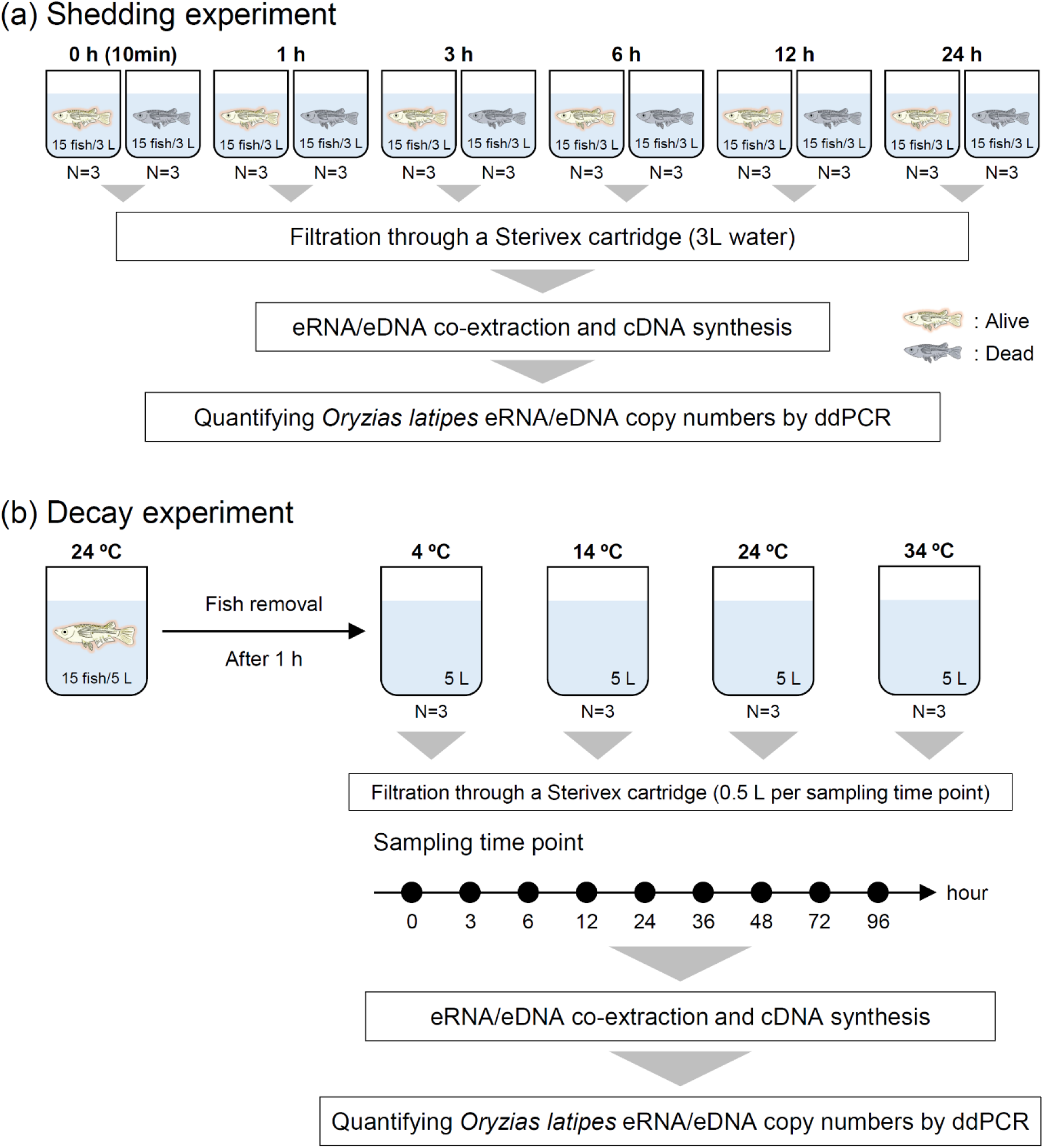
Experimental design. (a) Shedding experiment for eRNA/eDNA from *Oryzias latipes*. Each aquarium (n = 3) contained 3 L of dechlorinated water with fifteen live or dead *Oryzias latipes* and was maintained at 24 ± 1 °C. The full 3 L water volume was filtered through 0.45-µm Sterivex cartridges at 0 (10 min), 1, 3, 6, 12, and 24 h. (b) Decay experiment for eRNA/eDNA from *O. latipes*. Each aquarium (n = 3) contained 5 L of dechlorinated water in which fifteen *O. latipes* were held for 1 h before removal. Aquaria were incubated at 4, 14, 24, or 34 °C, and 0.5 L water samples were collected up to 96 h and filtered through 0.45-µm Sterivex cartridges. Co-extraction of eRNA/eDNA, purification, cDNA synthesis, and digital polymerase chain reaction (ddPCR) quantification were performed to estimate eNA shedding and decay dynamics.

At each time point, the full 3 L water volume was filtered through a Sterivex cartridge (0.45 µm; Millipore SVHV010RS; Merck, Tokyo, Japan) at 0 (10 min), 1, 3, 6, 12, and 24 h. Sterivex filters were stored at 4 °C, and eRNA/eDNA were extracted as soon as possible; if extraction was delayed by more than 6 h, filters were stored at −20 °C. Water temperature, pH, and dissolved oxygen (DO) were measured at each sampling time point using a digital thermometer SN3000 (NETSUKEN Corp., Tokyo, Japan) and a pH/DO meter D-55 (HORIBA, Kyoto, Japan). A parallel experiment quantified eRNA/eDNA release from dead *O. latipes* using euthanized fish instead of live individuals. Fish were euthanized by immersion in ice water for 1 h before being introduced into the aquaria.

### eRNA/eDNA decay from *Oryzias latipes*

Glass aquaria were filled with 5 L of dechlorinated water, and the temperature was maintained at 24 ± 1 °C using an open-circuit chiller (Taitec Corp., Saitama, Japan). Fifteen live fish were placed in each aquarium for 1 h under continuous aeration and then removed (Figure 1b). Aquaria were then incubated at 4, 14, 24, and 34 °C in temperature-controlled water baths; the time of placement was defined as time 0 (*t* = 0). We prepared triplicate aquaria for each temperature, along with a negative control aquarium containing dechlorinated water without fish. At each temperature, 0.5 L water samples were collected and filtered through Sterivex cartridges (0.45 µm; Millipore SVHV010RS; Merck, Tokyo, Japan) at 0, 3, 6, 12, 24, 36, 48, 72, and 96 h. Sterivex filters were stored at 4 °C, and eRNA/eDNA were extracted as soon as possible; if extraction was delayed by more than 6 h, filters were stored at −20 °C.

The shedding and decay experiments followed the guidelines of the Experimental Animal Facility of Kao Corporation’s R&D Department.

### eRNA/eDNA co-extraction and cDNA synthesis

Total DNA/RNA was co-extracted using the AllPrep DNA/RNA Mini Kit (Qiagen, Hilden, Germany). Buffer RTL Plus (1 mL) was introduced into each Sterivex cartridge through the inlet, and the cartridges were incubated with moderate rotation at 60 °C for 60 min. The lysis buffer was collected by centrifuging each cartridge in a 50-mL tube at 8000 × g for 1 min and then transferred to an AllPrep DNA spin column. Subsequent steps followed the manufacturer’s protocol, with a 50 μL elution volume. DNA samples were purified using a NucleoSpin DNA Clean-up XS kit (MACHEREY-NAGEL, Düren, Germany). Before cDNA synthesis, the RNA samples underwent two consecutive DNase treatments with an rDNase set and were purified using a NucleoSpin RNA Clean-up XS kit (MACHEREY-NAGEL, Düren, Germany). cDNA was synthesized from single-stranded RNA using a PrimeScript II 1^st^ Strand cDNA Synthesis Kit (TaKaRa Bio Inc., Shiga, Japan) according to the manufacturer’s protocol. To assess cross-contamination during sample filtration, DNA/RNA co-extraction, and cDNA synthesis, we simultaneously extracted RNA and DNA from deionized water as negative controls. All extracted DNA/RNA and cDNA samples were immediately stored at −80 °C.

### Quantifying medaka eDNA/eRNA and microbial DNA by ddPCR

eRNA/eDNA in each water sample was quantified using a Bio-Rad QX200 AutoDG Droplet Digital PCR system (Bio-Rad Laboratories, Hercules, CA, USA), with negative controls for filtration, co-extraction, and cDNA synthesis. Before digital polymerase chain reaction (ddPCR) analysis, DNA and cDNA were diluted to reduce inhibition. PCR was performed in a 22-μL reaction volume containing 11 μL of 2× ddPCR Supermix for probe, 1.1 μL template DNA or cDNA, 2.0 μL of NIES_R primer mix (forward and reverse primers, 10 µM), 0.6 μL NIES_R probe (Supplementary Table S1), and 5.3 μL sterile deionized water. Thermal cycling consisted of 95 °C for 5 min; 40 cycles of 94 °C for 30 s and 54 °C for 1 min; followed by 4 °C for 5 min and 90 °C for 5 min. Microbial DNA was quantified by ddPCR with EvaGreen. PCR was conducted in a 22-μL reaction containing 11 μL of 2× ddPCR Supermix for EvaGreen, 1.1 μL template DNA, 0.3 μL primer mix (341F and R806, 10 μM; Supplementary Table S1), and 9.3 μL sterile deionized water. Thermal cycling comprised 95 °C for 5 min; 50 cycles of 95 °C for 30 s and 55 °C for 2 min; followed by 4 °C for 5 min and 95 °C for 5 min.

Each sample was quantified in triplicate, with negative controls (1.1 μL sterile distilled water) and positive controls included in each plate. The positive control comprised 1.1 μL total DNA extracted from *O. latipes* or activated sludge. The DNA concentration of the positive control was 0.2 ng/μL, measured using a Qubit 4.0 Fluorometer (Invitrogen, Carlsbad, CA, USA). Data were analyzed with Bio-Rad QuantaSoft software version 1.7.4 (Bio-Rad Laboratories). Samples with fewer than 10,000 accepted droplets were excluded and re-measured until at least 10,000 accepted droplets were obtained. The droplet-positivity threshold was set manually. The limit of detection (LOD) was defined as 0.1 copies/μL, following previous studies^18,27,28^. eRNA/eDNA copy numbers were calculated as the mean of triplicate measurements. To convert ddPCR concentrations to copy numbers per original water volume filtered, we applied Equation (1):

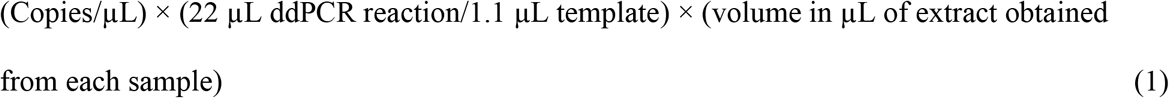

### Degradation kinetic analysis

The eNA decay rate was modeled using a first-order exponential decay model as represented by Equation (2):^27,29,30^

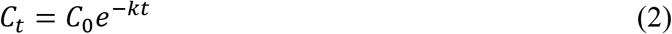

where *C_t_* is the eNA concentration at time *t*, *C_0_* is the eNA concentration at *t* = 0, and *k* is the first-order decay rate constant (h^−1^).

The half-life (*t_1/2_*) (h), defined as the time required for *C_t_* to decrease to 50% of *C_0_*, was calculated using Equation (3):^31^

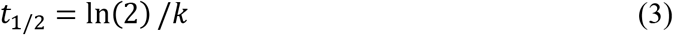

Temperature dependence of *k* was described by the Arrhenius equation (Equation (4)):^22,31^

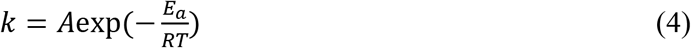

where *E_a_* is the activation energy (kJ mol^−1^), *A* is the frequency factor, *R* is the universal gas constant (8.314 Jmol^−1^K^−1^), and *T* is the absolute temperature (K). *Ea* was estimated from the slope of Arrhenius plots of ln(*k*) versus 1/*T*, as represented by Equation (5):

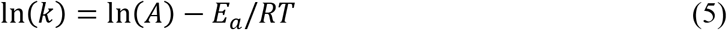

The temperature coefficient (*Q_10_*) was calculated using Equation (6): ^24^

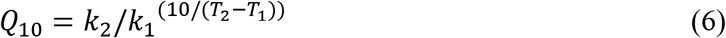

where *k_1_* and *k_2_* are decay rate constants at the temperature *T_1_* and *T_2_* (°C), respectively.

The decimal reduction time (*D* value) and the thermal resistance coefficient *(z* value) of eNA decay were calculated using Equations (7) and (8):^32,33^

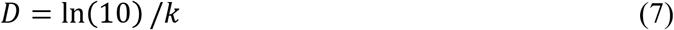

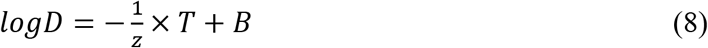

where *T* is the temperature (°C), and *B* is the linear coefficient. The *z* value was estimated from the slope of log*D* versus *T*.

### Statistical analysis

All statistical analyses were performed in Origin Pro 2021b (OriginLab, Northampton, MA, USA) or R version 4.3.1 (R Core Team, Vienna, Austria), with significance set at *p* = 0.05. Normality was assessed using the Shapiro–Wilk test. Differences in eRNA/eDNA copy numbers released from live and dead *O. latipes* were evaluated using Welch’s *t*-test with Holm correction because the datasets were normally distributed. First-order decay rate constants (*k*) were estimated by fitting an exponential decay model (Equation 2) to raw time-series data. Differences in *k* between eRNA and eDNA were assessed using a linear mixed-effects model generated with the R package lme4. The model used log_e_-transformed copy number (copied/0.5 L) of eDNA and eRNA as the response variable; sampling time (h), eNA type (eDNA or eRNA), temperature (4, 14, 24, and 34 °C), and the three-way interaction among sampling time, eNA type, and temperature as fixed effects; and glass-tank replicate as a random effect. Post hoc Holm tests performed with the R package emmeans were used to evaluate differences in decay constants between eNA types.

## RESULTS AND DISCUSSION

### Medaka measurements and physicochemical conditions in shedding and decay experiments

The mean (±SD) length and mass of *O. latipes* used in the experiments were 2.64 ± 0.139 cm and 0.202 ± 0.0315 g, respectively. In the shedding experiment, pH and temperature remained relatively stable in each aquarium, whereas DO declined over time despite aeration (Supplementary Table S2). In the aquarium containing dead *O. latipes*, DO decreased to 0.45 mg/L after 24 h. In the decay experiment, pH, DO, and temperature remained relatively constant in all aquaria (Supplementary Table S3). All negative controls, including fish-free aquaria, filter controls, extraction/purification/cDNA synthesis controls, and PCR negative controls, remained below the LOD throughout both experiments.

### Quantification of eRNA/eDNA shedding from live and dead *O. latipes*

In the shedding experiment, medaka-derived eRNA and eDNA exhibited distinct temporal patterns in aquaria containing live and dead *O. latipes* (Figure 2a, b). In both treatments, eDNA release was higher than eRNA release. Mean eRNA copy numbers peaked at 3 h in the dead-fish aquaria and at 6 h in the live-fish aquaria, whereas mean eDNA copy numbers peaked at 12 h in both treatments. After these peaks, eRNA and eDNA copy numbers declined markedly over time. In contrast, microbial DNA copy numbers increased approximately logarithmically throughout the observation period, consistent with rapid microbial proliferation (Supplementary Figure S1). These patterns suggest that eNA release from live or dead fish and microbial eRNA/eDNA degradation processes may have occurred simultaneously. After *O. latipes*-derived eNA peaked, degradation likely became dominant relative to continued release. The earlier eRNA peak in the dead-fish treatment may reflect faster eRNA degradation under heightened microbial activity during medaka decomposition. The sharp DO decline observed only in dead-fish aquaria at 24 h further supports increased microbial activity.^27,34^

**Figure 2.**
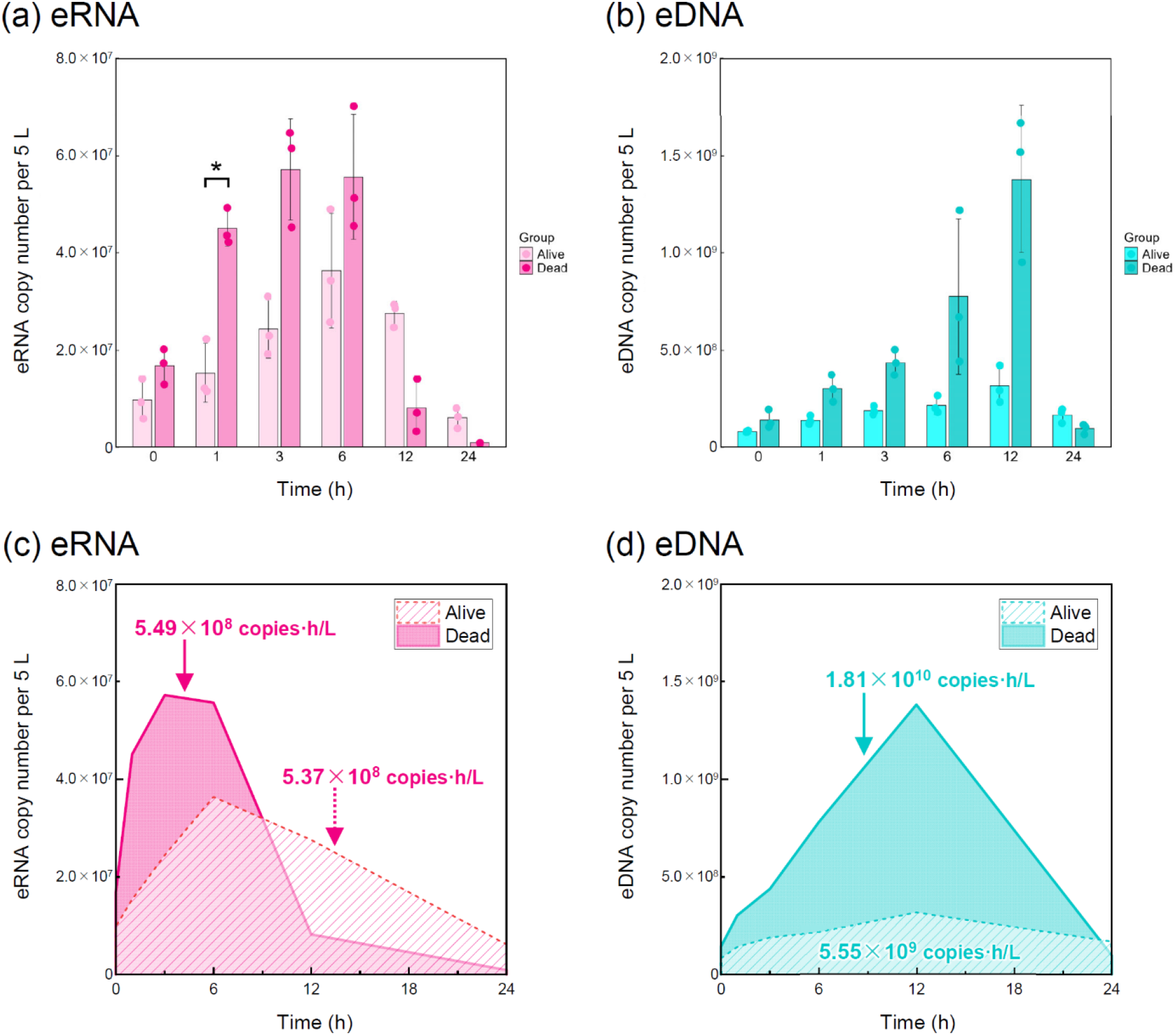
Comparison of eRNA/eDNA copy numbers from live and dead *Oryzias latipes*. Bar graph shows mean copy numbers of (a) eRNA and (b) eDNA in aquaria containing live and dead *O. latipes*; points indicate three replicates. Error bars represent standard deviation (SD). Differences in eRNA/DNA shedding between live and dead *O. latipes* were analyzed using Welch’s *t*-test. Asterisks indicate statistically significant differences after Holm correction (\**p* < 0.05). Note: *y*-axis scales differ between (a) and (b). Area under the curve (AUC) was calculated from time-series profiles of mean (c) eRNA and (d) eDNA copy numbers in the shedding experiment.

Welch’s *t*-test showed that mean eRNA copy numbers from dead *O. latipes* were significantly higher than those from live fish only at 1 h (*p* = 0.023). No significant differences were detected at other time points; however, eRNA release from dead medaka exceeded that from live medaka through 6 h, and eDNA release from dead medaka exceeded that from live medaka through 12 h. To compare cumulative shedding between treatments, we calculated the area under the curve (AUC) for the 0–24 h time-series profiles (copies·h/L; trapezoidal integration; Figure 2c, d). For eRNA, the AUC was slightly higher in the dead-fish treatment (5.49 × 10^8^ copies·h/L) than in the live-fish treatment (5.37 × 10^8^ copies·h/L). For eDNA, the dead-fish AUC (1.81×10^10^ copies·h/L) was approximately three times higher than the live-fish AUC (5.55 × 10^9^ copies·h/L), suggesting that carcasses may contribute more total eNA than live individuals and may release proportionally more eDNA than eRNA. In natural environments, eDNA-based analyses may therefore retain detectable carcass-derived signals and continue to detect organisms after they are no longer locally present. This interpretation aligns with Miyata et al. and with studies showing that eNA transport and migratory fish movement can generate detections of taxa absent from the sampling location at the time of collection.^4,11,13,35^ Together, these findings support the potential value of eRNA as a marker for more recent organismal presence. Further field-based studies are warranted to clarify the environmental fate, transport, and persistence of carcass-derived eNA under realistic conditions.

We hypothesized that dead *O. latipes* would not release detectable eRNA. Consistent with previous reports, our time-series measurements showed greater eDNA release from dead fish than from live *O. latipes*, suggesting that carcass-derived eDNA originated from tissues and cells sloughed off during decomposition.^14^ However, dead *O. latipes* also produced detectable eRNA. Given that the dead fish treatment used freshly euthanized individuals, the detected eRNA may have been protected within cells associated with scales, mucus, feces, and tissue fragments released from fresh carcasses. Additionally, Pozhitkov et al. reported that many genes associated with stress responses, immunity, inflammation, and apoptosis can persist, and in some cases increase after death in mice and zebrafish.^36^ In this context, eRNA detected in the dead-fish aquaria may reflect both RNA retained within cellular material and post-mortem transcript dynamics. Hence, carcasses can be substantial sources of eRNA and eDNA shortly after death, and eRNA analysis does not necessarily exclude carcass-derived signals. However, the controlled time-course patterns provide important evidence supporting the utility of eRNA monitoring: mean eRNA copy numbers in the live-fish treatment exceeded those in the dead-fish treatment at 12 h, whereas mean eDNA copy numbers did not exceed those in the dead-fish treatment until 24 h (Figure 2). Thus, when carcasses are near a sampling site, eDNA-based surveys may remain affected by carcass-derived signals for at least 24 h, while eRNA-based surveys may become less affected within 12 h of a mortality event, potentially improving inference of signals more closely associated with living individuals.

Overall, these findings suggest that carcass-derived eNA can lead to false inference that a target species is currently alive at a sampling site when carcasses are present. This risk is particularly relevant for eDNA, which is generally more persistent and may therefore provide longer-lasting false-positive signals. To minimize carcass-derived signals, eRNA analysis may be useful as eRNA can be released in lower absolute amounts and degrade more rapidly than eDNA; complementary approaches such as the eDNA integrity index (eDI), which accounts for DNA fragment length, may also help distinguish recent from older signals. ^37–39^

### Temperature-dependent differences in eRNA and eDNA decay rates

In the decay experiment, eRNA and eDNA concentrations declined over time and were well described by first-order exponential decay models (Figure 3), with coefficients of determination (*R*^2^) ranging from 0.75 to 0.96 across temperature treatments. Estimated decay rate constants (*k,* h^−1^) for eRNA and eDNA were 0.0268 and 0.0179 at 4 °C, 0.0368 and 0.0281 at 14 °C, 0.128 and 0.0367 at 24 °C, and 0.182 and 0.0427 at 34 °C, respectively. eRNA decay rates were significantly higher than eDNA decay rates at 14, 24, and 34 °C (14 °C: *p* = 0.0116, 24 and 34 °C: *p* < 0.0001), but not at 4 °C (Supplementary Table S4). Half-lives calculated from *k* values were 25.8, 18.9, 5.40, and 3.81 h for eRNA at 4, 14, 24, and 34 °C, respectively, compared with 38.6, 24.7, 18.9, and 16.2 h for eDNA (Table 1). Thus, eRNA had shorter half-lives than eDNA across all temperature treatments. As enzyme kinetics and microbial growth generally increase with temperature up to an optimum,^40^ the faster increase in microbial abundance observed at higher temperatures suggests that microbial activity may have contributed to eNA degradation in the aquaria (Supplementary Figure S2). Moreover, DNA and RNA hydrolysis accelerates as temperature rises.^41–43^ Together, these processes plausibly explain the observed temperature-dependent increase in eNA decay rates.

**Figure 3.**
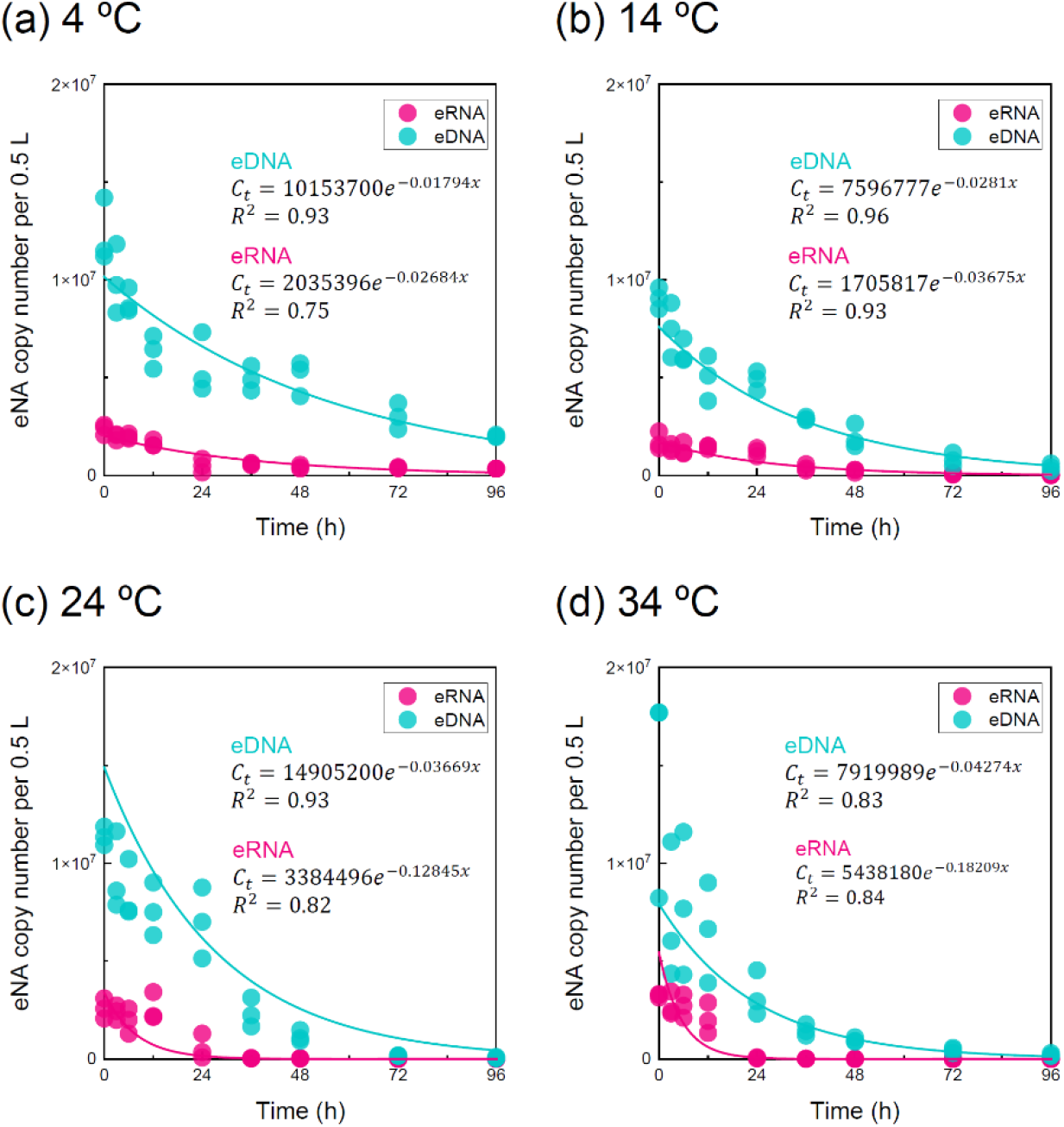
Time-dependent changes in eRNA/eDNA copy numbers for (a) 4, (b) 14, (c) 24, and (d) 34 °C in triplicate experiments. Curves were fitted using a first-order exponential decay model, and *R*^2^ values indicate the model’s goodness of fit.

**Table 1.** Decay rate constants and half-life time for *Oryzias latipes* eRNA/eDNA in the aquarium experiment.

|  | Temperature (°C) | Decay constant (/h) | Half-life (h) |
| --- | --- | --- | --- |
| eRNA | 4 | 0.0268 | 25.8 |
|  | 14 | 0.0367 | 18.9 |
|  | 24 | 0.128 | 5.40 |
|  | 34 | 0.182 | 3.81 |
| eDNA | 4 | 0.0179 | 38.6 |
|  | 14 | 0.0281 | 24.7 |
|  | 24 | 0.0367 | 18.9 |
|  | 34 | 0.0427 | 16.2 |

Previous studies have reported contrasting temperature effects on eRNA and eDNA persistence. Qian et al. found that prawn eRNA decays faster at lower temperatures, whereas eDNA decays faster than eRNA at higher temperatures.^20^ In contrast, Jo et al. reported that zebrafish eRNA decays faster than eDNA and that both decay more rapidly with warming, consistent with our findings.^21^ These discrepancies may reflect taxon-specific variation in how eNA is released and protected in the environment, such as associations with epithelial or sloughed cells, mucus, feces, and tissue fragments in fish versus exoskeleton-derived material in crustaceans.^18–21,44^ Similar release forms in small fish, including medaka and zebrafish, may explain their comparable decay patterns.^45–47^ Broader cross-taxon datasets spanning wider temperature ranges are needed to generalize eRNA/eDNA persistence in natural systems.

These observations suggest that eRNA may provide stronger temporal resolution than eDNA for detecting recent organismal presence during warm seasons. However, at 4 °C, eRNA and eDNA decay rates were comparable, indicating that eRNA is not necessarily short-lived under cold conditions. During colder periods, eRNA detections may persist longer than expected, potentially reducing the extent to which eRNA lowers false-positive risk relative to eDNA.

### Arrhenius-based interpretation of temperature-dependent eRNA/eDNA degradation

To compare the thermal sensitivity of eRNA and eDNA decay beyond single-temperature rate constants and half-lives, we calculated Arrhenius-based metrics (*Q_10_*, *E_a_*, and *z*-values; Table 2). *Q_10_*, the ratio of reaction rates associated with a 10 °C temperature increase, was calculated for three intervals: 4–14, 14–24, and 24–34 °C. For eDNA, *Q_10_* values were 1.57, 1.31, and 1.16, respectively, indicating a gradual increase in degradation rate with temperature. For eRNA, corresponding values were 1.37, 3.50, and 1.42, showing particularly strong thermal sensitivity between 14 and 24 °C and a non-linear temperature response across the full range.

**Table 2.**
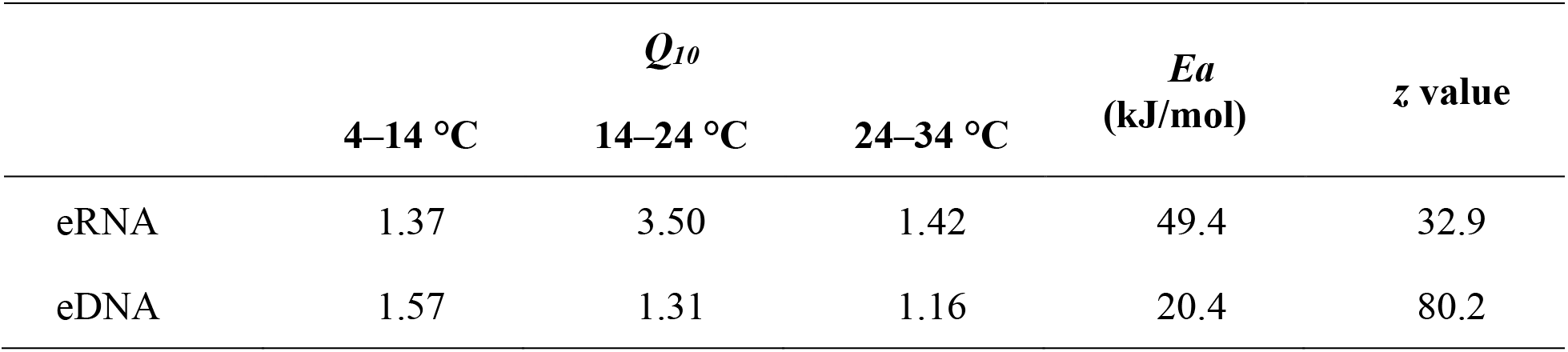
Temperature-dependence metrics (*Q*_10_, *E_a_*, and *z* values) for eRNA and eDNA estimated from decay experiments at four temperatures.

| | $Q_{10}$ | | | $E_a$<br>(kJ/mol) | $z$ value |
| --- | --- | --- | --- | --- | --- |
|  | 4–14 °C | 14–24 °C | 24–34 °C |  |  |
| eRNA | 1.37 | 3.50 | 1.42 | 49.4 | 32.9 |
| eDNA | 1.57 | 1.31 | 1.16 | 20.4 | 80.2 |

Activation energy (*E_a_*), derived from the slope of the Arrhenius plot, provides an index of the temperature dependence; higher values indicate stronger dependence. The *E_a_* for eRNA (49.4 kJ mol^−1^) was approximately twice that for eDNA (20.4 kJ mol^−1^), suggesting that eRNA decay was more temperature-dependent (Figure 4, Table 2).

**Figure 4.**
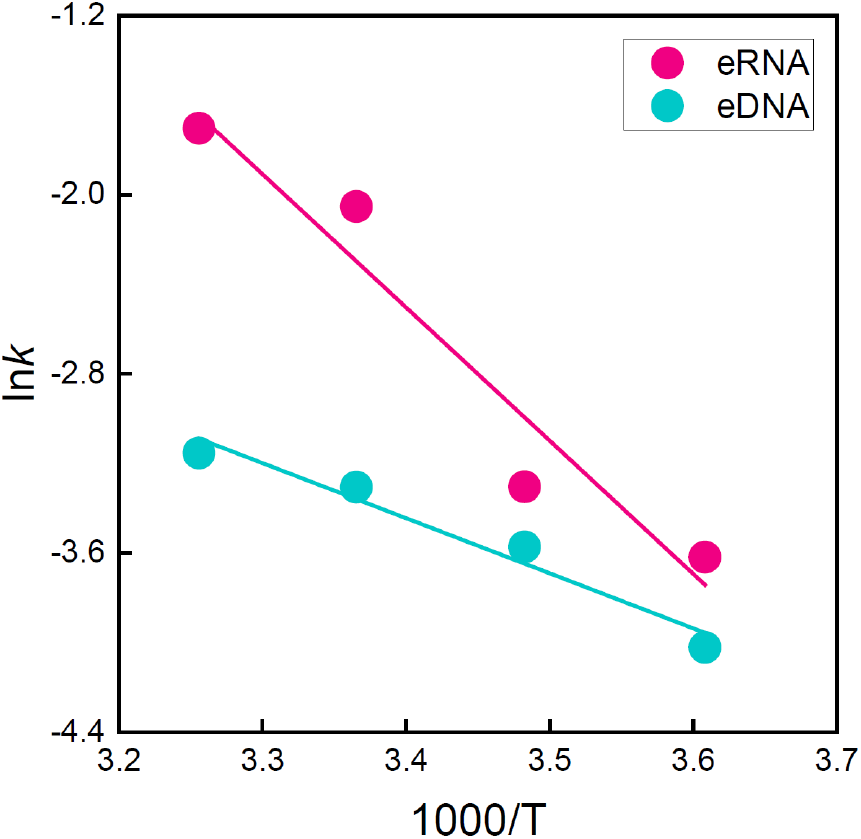
Arrhenius plot for eRNA and eDNA decay rate constants. ln(*k*) is plotted against 1/*T* (K^−1^) for eRNA and eDNA; slopes were used to estimate activation energies (*E_a_*). Coefficients of determination (*R*^2^) for the linear model are 0.90 for eRNA and 0.94 for eDNA.

For a single chemical reaction, activation energy represents the minimum energy required for the reaction to proceed. In this study, however, it reflects an apparent activation energy arising from multiple simultaneous processes, including microbial uptake, disruption of protective structures surrounding eNAs, and enzymatic degradation. Although the mechanisms underlying the different *E_a_* values for eRNA and eDNA cannot be identified definitively, the higher eRNA value may reflect differences in the optimal temperature ranges of eRNA- and eDNA-degrading enzymes and in the environmental forms of the two molecules, with eRNA potentially more strongly protected within cells, extracellular vesicles, and protein capsids.^48^

The *z*-value, another indicator of temperature sensitivity, was lower for eRNA (32.9) than for eDNA (80.2), consistent with greater thermal sensitivity of eRNA decay; lower *z*-values combined with higher *E_a_* values generally indicate stronger temperature dependence.^33,49^ Together, these metrics demonstrate that eRNA and eDNA degradation differ substantially in thermal sensitivity. Given that eRNA degrades rapidly under warm conditions, eRNA-based biomonitoring may provide higher temporal resolution for recent organismal presence than eDNA. Under cold conditions, however, eRNA degradation rates may be comparable to those of eDNA; thus, eRNA detection alone may not provide unambiguous evidence of living organisms. Nevertheless, eRNA-based monitoring should not be considered inherently disadvantageous in cold periods. Several studies have reported fewer false positives with eRNA than with eDNA even at low temperatures, suggesting that factors other than temperature can strongly influence eNA persistence in natural environments.^11,50^ Additional field studies are needed to evaluate the utility of eRNA analysis during cold periods.

This study has certain limitations that should be considered. First, as it was conducted in controlled aquarium systems and relied on a single model fish species and a single mitochondrial marker, the findings should be generalized with caution to natural environments and other taxa.^44^ Natural systems include additional processes not represented here, including variation in water chemistry, UV exposure, sediment interactions, and more complex microbial communities, all of which may influence eNA shedding and decay.^27^ Future studies should evaluate carcass-derived eRNA/eDNA signals across broader taxonomic groups and genetic markers under more realistic environmental conditions. Second, because this study used freshly euthanized *O. latipes,* future work should examine how decomposition stage affects eNA release and carcasses’ contribution to environmental signals. Finally, direct comparisons between eRNA-based analyses and conventional capture-based surveys during cold periods will be important for developing practical field-monitoring guidance.

## CONCLUSION

We quantified carcass-derived environmental nucleic acid (eNA) release and assessed the temperature-dependent decay of eRNA and eDNA to clarify how carcasses may affect eNA-based monitoring in natural environments. *O. latipes* carcasses released detectable eDNA and eRNA, indicating that eRNA-based biomonitoring does not necessarily eliminate carcass-derived signals. Arrhenius-based kinetic parameters showed clear differences in temperature dependence between eRNA and eDNA. At lower temperatures, eRNA and eDNA decay rates were comparable, whereas eRNA decayed significantly faster than eDNA at higher temperatures. These findings suggest that eRNA-based biomonitoring may improve temporal resolution for detecting recent presence under warm conditions, but its relative advantage may decrease under cold conditions. For eNA-based biomonitoring in natural environments, we recommend: (1) when carcasses are observed, record their species identity and location and incorporate this information into data interpretation; (2) to mitigate false-positive risk, use eRNA, particularly under warm conditions where it is expected to degrade rapidly, and/or complementary metrics such as the eDI; and (3) adopt temperature-aware sampling strategies, including increased temporal replication during cold periods. Together, these measures can reduce false-positive risk and strengthen inference reliability in aquatic biomonitoring.

## ASSOCIATED CONTENT

### Supporting Information

The Supporting Information is available free of charge at URL. Time-dependent change in bacterial 16S rRNA gene copy number, primer pairs used in this study, water quality parameters in experiments, and summary of post-hoc Holm test results from a linear mixed model comparing decay rate constants (PDF)Time-dependent change in bacterial 16S rRNA gene copy number, primer pairs used in this study, water quality parameters in experiments, and summary of post-hoc Holm test results from a linear mixed model comparing decay rate constants (PDF)

## AUTHOR INFORMATION

### Author Contributions

Conceptualization, Y.I.; K.M. Methodology, Y.I.; K.M.; T.M. Formal Analysis, Y.I. Visualization, Y.I. Investigation, Y.I.; K.M.; H.H. Project Administration, Y.I.; K.M. Writing–Original Draft Preparation, Y.I. Writing–Review and Editing, Y.I, K.M., T.M., H.H., and M.Y. Supervision, H.H.; M.Y.

### Conflict of Interest Statement

The authors have no conflicts of interest to declare

## ACKNOWLEDGMENT

We thank Mrs. Tomoko Yokomatsu for her support in managing the Japanese medaka breeding system and maintaining the aquarium.

## SYNOPSIS

Fish carcasses release detectable eRNA, and temperature governs its interpretability: eRNA can improve “recent presence” resolution in warm water but may persist long enough in cold water to capture carcass signals and trigger false positives.

## TABLE OF CONTENTS

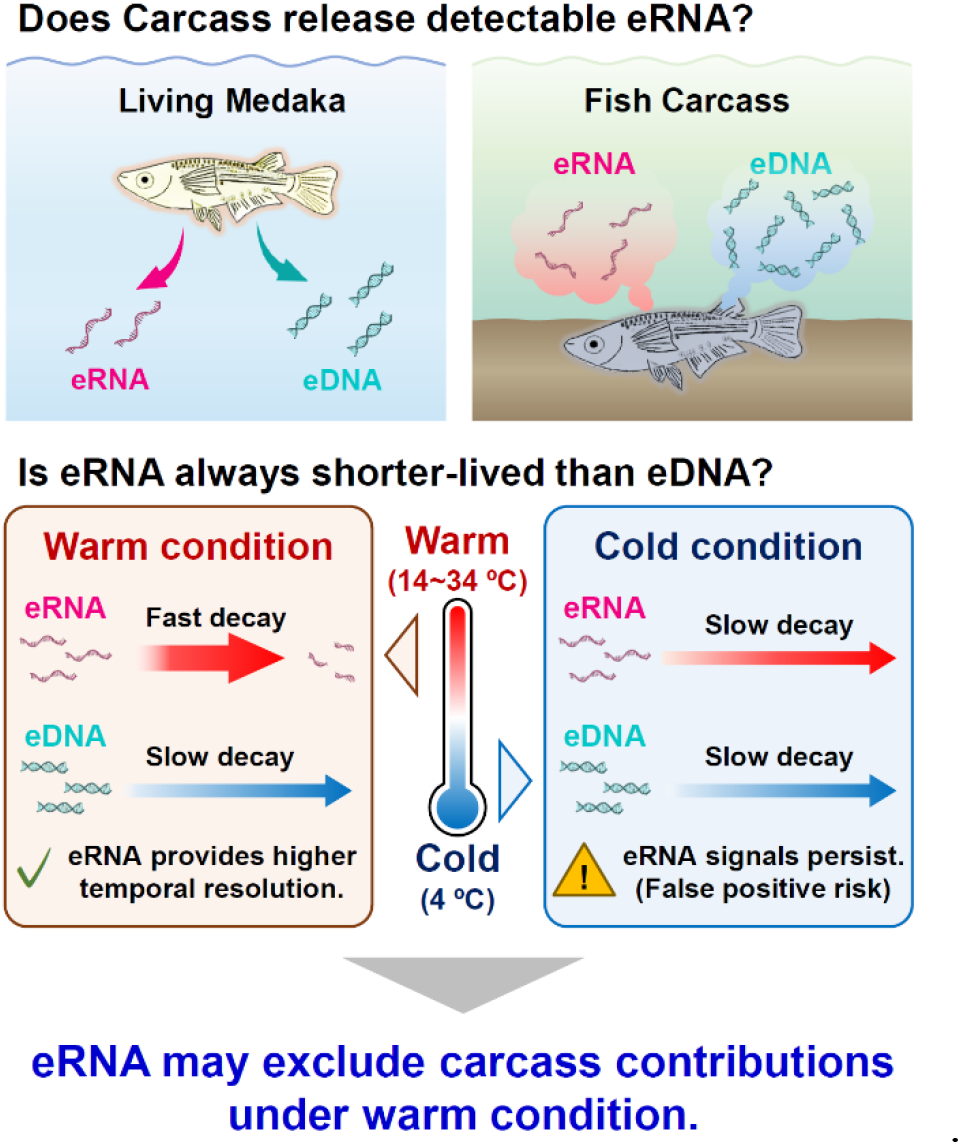

## Notes

### Competing Interest Statement

The authors have declared no competing interest.

## REFERENCES

(1) Taberlet, P.; Coissac, E.; Hajibabaei, M.; Rieseberg, L. H. Environmental DNA. Mol. Ecol. 2012, 21 (8), 1789–1793. 10.1111/j.1365-294X.2012.05542.x

(2) Thomsen, P. F.; Willerslev, E. Environmental DNA – An Emerging Tool in Conservation for Monitoring Past and Present Biodiversity. Biol. Conserv. 2015, 183, 4–18. 10.1016/j.biocon.2014.11.019

(3) Goldberg, C. S.; Turner, C. R.; Deiner, K.; Klymus, K. E.; Thomsen, P. F.; Murphy, M. A.; Spear, S. F.; McKee, A.; Oyler-McCance, S. J.; Cornman, R. S.; Laramie, M. B.; Mahon, A. R.; Lance, R. F.; Pilliod, D. S.; Strickler, K. M.; Waits, L. P.; Fremier, A. K.; Takahara, T.; Herder, J. E.; Taberlet, P. Critical Considerations for the Application of Environmental DNA Methods to Detect Aquatic Species. Methods Ecol. Evol. 2016, 7 (11), 1299–1307. 10.1111/2041-210X.12595

(4) Deiner, K.; Bik, H. M.; Mächler, E.; Seymour, M.; Lacoursière-Roussel, A.; Altermatt, F.; Creer, S.; Bista, I.; Lodge, D. M.; de Vere, N.; Pfrender, M. E.; Bernatchez, L. Environmental DNA Metabarcoding: Transforming How We Survey Animal and Plant Communities. Mol. Ecol. 2017, 26 (21), 5872–5895. 10.1111/mec.14350

(5) Turner, C. R.; Uy, K. L.; Everhart, R. C. Fish Environmental DNA Is More Concentrated in Aquatic Sediments than Surface Water. Biol. Conserv. 2015, 183, 93–102. 10.1016/j.biocon.2014.11.017

(6) Lee, H.-T.; Liao, C.-H.; Hsu, T.-H. Environmental DNA (eDNA) Metabarcoding in the Fish Market and Nearby Seafood Restaurants in Taiwan Reveals the Underestimation of Fish Species Diversity in Seafood. Biology 2021, 10 (11). 10.3390/biology10111132

(7) Inoue, Y.; Miyata, K.; Yamane, M.; Honda, H. Environmental Nucleic Acid Pollution: Characterization of Wastewater Generating False Positives in Molecular Ecological Surveys. ACS EST Water 2023. 10.1021/acsestwater.2c00542

(8) Li, Y.; Breaker, R. R. Kinetics of RNA Degradation by Specific Base Catalysis of Transesterification Involving the 2‘-Hydroxyl Group. J. Am. Chem. Soc. 1999, 121 (23), 5364–5372. 10.1021/ja990592p

(9) Chomczynski, P. Solubilzation in Formamide Protects RNA from Degradation. Nucleic Acids Res. 1992, 20 (14), 3791–3792. 10.1093/nar/20.14.3791

(10) Cristescu, M. E. Can Environmental RNA Revolutionize Biodiversity Science? Trends Ecol. Evol. 2019, 34 (8), 694–697. 10.1016/j.tree.2019.05.003

(11) Miyata, K.; Inoue, Y.; Amano, Y.; Nishioka, T.; Yamane, M.; Kawaguchi, T.; Morita, O.; Honda, H. Fish Environmental RNA Enables Precise Ecological Surveys with High Positive Predictivity. Ecol. Indic. 2021, 128, 107796. 10.1016/j.ecolind.2021.107796

(12) Miyata, K.; Inoue, Y.; Amano, Y.; Nishioka, T.; Nagaike, T.; Kawaguchi, T.; Morita, O.; Yamane, M.; Honda, H. Comparative Environmental RNA and DNA Metabarcoding Analysis of River Algae and Arthropods for Ecological Surveys and Water Quality Assessment. Sci. Rep. 2022, 12 (1), 19828. 10.1038/s41598-022-23888-1

(13) Miyata, K.; Kusakabe, Y.; Inoue, Y.; Yamane, M.; Honda, H. Validation of Fish Environmental RNA Metabarcoding Analysis for Ecological Surveys by Additional Traditional Field Surveys in the Naka River. Front. Ecol. Evol. 2025, 13. 10.3389/fevo.2025.1540001

(14) Tillotson, M. D.; Kelly, R. P.; Duda, J. J.; Hoy, M.; Kralj, J.; Quinn, T. P. Concentrations of Environmental DNA (eDNA) Reflect Spawning Salmon Abundance at Fine Spatial and Temporal Scales. Biol. Conserv. 2018, 220, 1–11. 10.1016/j.biocon.2018.01.030

(15) Payne, J. A. A Summer Carrion Study of the Baby Pig Sus Scrofa Linnaeus. Ecology 1965, 46 (5), 592–602. 10.2307/1934999

(16) Merkes, C. M.; McCalla, S. G.; Jensen, N. R.; Gaikowski, M. P.; Amberg, J. J. Persistence of DNA in Carcasses, Slime and Avian Feces May Affect Interpretation of Environmental DNA Data. PLOS ONE 2014, 9 (11), e113346. 10.1371/journal.pone.0113346

(17) Dunker, K. J.; Sepulveda, A. J.; Massengill, R. L.; Olsen, J. B.; Russ, O. L.; Wenburg, J. K.; Antonovich, A. Potential of Environmental DNA to Evaluate Northern Pike (Esox Lucius) Eradication Efforts: An Experimental Test and Case Study. PLOS ONE 2016, 11 (9), e0162277. 10.1371/journal.pone.0162277

(18) Wood, S. A.; Biessy, L.; Latchford, J. L.; Zaiko, A.; von Ammon, U.; Audrezet, F.; Cristescu, M. E.; Pochon, X. Release and Degradation of Environmental DNA and RNA in a Marine System. Sci. Total Environ. 2020, 704, 135314. 10.1016/j.scitotenv.2019.135314

(19) Marshall, N. T.; Vanderploeg, H. A.; Chaganti, S. R. Environmental (e)RNA Advances the Reliability of eDNA by Predicting Its Age. Sci. Rep. 2021, 11 (1), 2769. 10.1038/s41598-021-82205-4

(20) Qian, T.; Shan, X.; Wang, W.; Jin, X. Effects of Temperature on the Timeliness of eDNA/eRNA: A Case Study of Fenneropenaeus Chinensis. Water 2022, 14 (7). 10.3390/w14071155

(21) Jo, T.; Tsuri, K.; Hirohara, T.; Yamanaka, H. Warm Temperature and Alkaline Conditions Accelerate Environmental RNA Degradation. Environ. DNA 2023, 5 (5), 836–848. 10.1002/edn3.334

(22) Arrhenius, S. Über die Reaktionsgeschwindigkeit bei der Inversion von Rohrzucker durch Säuren. Z. Phys. Chem. 1889, 4U (1), 226–248. 10.1515/zpch-1889-0416

(23) Leahy, J. G.; Colwell, R. R. Microbial Degradation of Hydrocarbons in the Environment. Microbiol. Rev. 1990, 54 (3), 305–315. 10.1128/mr.54.3.305-315.1990

(24) Gillooly, J. F.; Brown, J. H.; West, G. B.; Savage, V. M.; Charnov, E. L. Effects of Size and Temperature on Metabolic Rate. Science 2001, 293 (5538), 2248–2251. 10.1126/science.1061967

(25) Brown, J. H.; Gillooly, J. F.; Allen, A. P.; Savage, V. M.; West, G. B. Toward a Metabolic Theory of Ecology. Ecology 2004, 85 (7), 1771–1789. 10.1890/03-9000

(26) Schulte, P. M. The Effects of Temperature on Aerobic Metabolism: Towards a Mechanistic Understanding of the Responses of Ectotherms to a Changing Environment. J. Exp. Biol. 2015, 218 (12), 1856–1866. 10.1242/jeb.118851

(27) Strickler, K. M.; Fremier, A. K.; Goldberg, C. S. Quantifying Effects of UV-B, Temperature, and pH on eDNA Degradation in Aquatic Microcosms. Biol. Conserv. 2015, 183, 85–92. 10.1016/j.biocon.2014.11.038

(28) Kagzi, K.; Hechler, R. M.; Fussmann, G. F.; Cristescu, M. E. Environmental RNA Degrades More Rapidly than Environmental DNA across a Broad Range of pH Conditions. Mol. Ecol. Resour. 2022, 22 (7), 2640–2650. 10.1111/1755-0998.13655

(29) Barnes, M. A.; Turner, C. R.; Jerde, C. L.; Renshaw, M. A.; Chadderton, W. L.; Lodge, D. M. Environmental Conditions Influence eDNA Persistence in Aquatic Systems. Environ. Sci. Technol. 2014, 48 (3), 1819–1827. 10.1021/es404734p

(30) Dejean, T.; Valentini, A.; Duparc, A.; Pellier-Cuit, S.; Pompanon, F.; Taberlet, P.; Miaud, C. Persistence of Environmental DNA in Freshwater Ecosystems. PLOS ONE 2011, 6 (8), e23398. 10.1371/journal.pone.0023398

(31) Atkins, P. W. Atkins’ Physical Chemistry., Twelfth edition / Peter Atkins, Julio de Paula, James Keeler.; University Press: Oxford, 2022.

(32) van Asselt, E. D.; Zwietering, M. H. A Systematic Approach to Determine Global Thermal Inactivation Parameters for Various Food Pathogens. Int. J. Food Microbiol. 2006, 107 (1), 73–82. 10.1016/j.ijfoodmicro.2005.08.014

(33) Kim, A.-N.; Kim, O. W.; Kim, H. Degradation Kinetics of Physicochemical and Sensory Properties of Rice during Storage at Different Temperatures. LWT 2022, 164, 113688. 10.1016/j.lwt.2022.113688

(34) Price, P. B.; Sowers, T. Temperature Dependence of Metabolic Rates for Microbial Growth, Maintenance, and Survival. Proc. Natl. Acad. Sci. 2004, 101 (13), 4631–4636. 10.1073/pnas.0400522101

(35) Burgess, B. T.; Chea, R.; Westfall, K. M.; Abbott, C. L.; Iacarella, J. C. Environmental DNA Reflects Differences in Freshwater Habitat Use Between Two Pacific Salmonids at Fine Temporal Scales. Environ. DNA 2026, 8 (1), e70231. 10.1002/edn3.70231

(36) Pozhitkov, A. E.; Neme, R.; Domazet-Lošo, T.; Leroux, B. G.; Soni, S.; Tautz, D.; Noble, P. A. Tracing the Dynamics of Gene Transcripts after Organismal Death. Open Biol. 2017, 7 (1), 160267. 10.1098/rsob.160267

(37) Jo, T.; Murakami, H.; Masuda, R.; Sakata, M. K.; Yamamoto, S.; Minamoto, T. Rapid Degradation of Longer DNA Fragments Enables the Improved Estimation of Distribution and Biomass Using Environmental DNA. Mol. Ecol. Resour. 2017, 17 (6), e25–e33. 10.1111/1755-0998.12685

(38) Jo, T.; Takao, K.; Minamoto, T. Linking the State of Environmental DNA to Its Application for Biomonitoring and Stock Assessment: Targeting Mitochondrial/Nuclear Genes, and Different DNA Fragment Lengths and Particle Sizes. Environ. DNA 2022, 4 (2), 271–283. 10.1002/edn3.253

(39) Ge, Q.; Piao, Y.; Li, Z.; Yang, Y.; Pan, M.; Bai, Y. Environmental DNA Integrity Index Is Sensitive for Species Biomass Estimation in Freshwater. Sci. Total Environ. 2025, 966, 178734. 10.1016/j.scitotenv.2025.178734

(40) Bello, M. D.; Abreu, C. I. Temperature Structuring of Microbial Communities on a Global Scale. Curr. Opin. Microbiol. 2024, 82, 102558. 10.1016/j.mib.2024.102558

(41) Lindahl, T.; Nyberg, B. Heat-Induced Deamination of Cytosine Residues in Deoxyribonucleic Acid. Biochemistry 1974, 13 (16), 3405–3410. 10.1021/bi00713a035

(42) Schoenmaker, L.; Witzigmann, D.; Kulkarni, J. A.; Verbeke, R.; Kersten, G.; Jiskoot, W.; Crommelin, D. J. A. mRNA-Lipid Nanoparticle COVID-19 Vaccines: Structure and Stability. Int. J. Pharm. 2021, 601, 120586. 10.1016/j.ijpharm.2021.120586

(43) Chheda, U.; Pradeepan, S.; Esposito, E.; Strezsak, S.; Fernandez-Delgado, O.; Kranz, J. Factors Affecting Stability of RNA – Temperature, Length, Concentration, pH, and Buffering Species. J. Pharm. Sci. 2024, 113 (2), 377–385. 10.1016/j.xphs.2023.11.023

(44) Che-Pelicier, A.; Hampton, H. G.; Sabadel, A. J. M.; Thomson Laing, G.; Miller, T.; Pochon, X. Release and Degradation of Environmental DNA and RNA From Eels in Aotearoa New Zealand. Environ. DNA 2025, 7 (3), e70128. 10.1002/edn3.70128

(45) Hiki, K.; Yamagishi, T.; Yamamoto, H. Environmental RNA as a Noninvasive Tool for Assessing Toxic Effects in Fish: A Proof-of-Concept Study Using Japanese Medaka Exposed to Pyrene. Environ. Sci. Technol. 2023, 57 (34), 12654–12662. 10.1021/acs.est.3c03737

(46) Miyata, K.; Inoue, Y.; Yamane, M.; Honda, H. Fish Environmental RNA Sequencing Sensitively Captures Accumulative Stress Responses through Short-Term Aquarium Sampling. Sci. Total Environ. 2025, 959, 178182. 10.1016/j.scitotenv.2024.178182

(47) Hiki, K.; Jo, T. S. Comprehensive Sequencing of Environmental RNA From Japanese Medaka at Various Size Fractions and Comparison With Skin Swab RNA. Environ. DNA 2025, 7 (3), e70137. 10.1002/edn3.70137

(48) Cristescu, M. E. Can Environmental RNA Revolutionize Biodiversity Science? Trends Ecol. Evol. 2019, 34 (8), 694–697. 10.1016/j.tree.2019.05.003

(49) Van Boekel, M. A. J. S. Kinetic Modeling of Food Quality: A Critical Review. Compr. Rev. Food Sci. Food Saf. 2008, 7 (1), 144–158. 10.1111/j.1541-4337.2007.00036.x

(50) Inoue, Y.; Miyata, K.; Yamane, M.; Honda, H. Environmental RNA/DNA Metabarcoding for Ecological Risk Assessment of Chemicals Based on Response of Benthic Communities in Natural Environments. bioRxiv June 1, 2025, p 2025.05.29.656937. 10.1101/2025.05.29.656937

