## Supporting Information for "Can carcasses cause environmental nucleic acid pollution? Implications for interpreting environmental RNA/DNA survey results"

### 1. Supplementary Figures

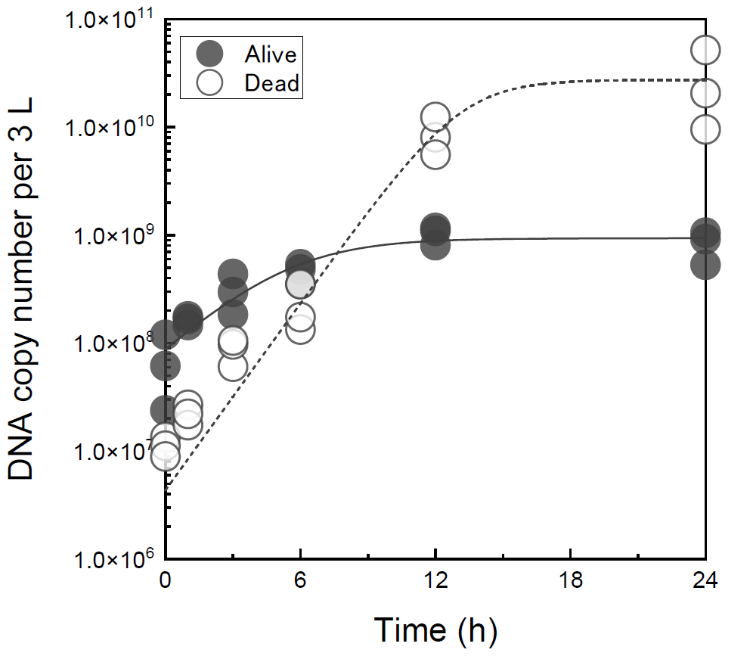


**Supplementary Figure S1.** Time-dependent changes in bacterial 16S rRNA gene copy number in aquaria containing live (●) and dead *Oryzias latipes* (○). The plots represent three replicates and lines represent the logistic model fits.

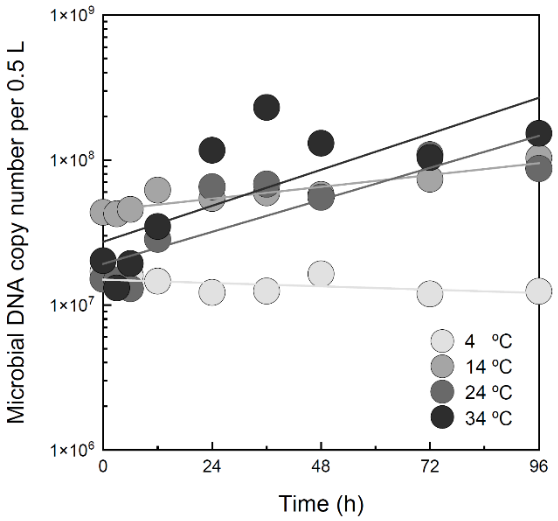


**Supplementary Figure S2.** Time-dependent changes in bacterial 16S rRNA gene copy number in decay experiments at 4, 14, 24, 34 °C. The plots represent the mean copy numbers of three replicates and lines represent the linear model fits.

### 2. Supplementary Tables

**Supplementary Table S1.** ddPCR primers and probes used to quantify Japanese medaka (*Oryzias latipes*) and bacterial 16S rRNA.

| **Species (target gene)** | **Primer name** | **Sequence (5' to 3')** | **Fluorophore/Quencher** |
|---|---|---|---|
| *Oryzias latipes* (*cytochrome b*) | NIES_R F | TTTGCCTACGCCATTCTACG | N/A |
| | NIES_R R | GGCTTCGTTGTTTAGAGGTGTG | N/A |
| | probe | TTAGCCTCTATTCTAGTACTATTC | FAM/QSY |
| Bacteria (*16S rRNA*) | 341F | CCTACGGGAGGCAGCAG | N/A |
| | R806 | GGACTACHVGGGTWTCTAAT | N/A |

**Supplementary Table S2.** Water quality parameters over time from live and dead *Oryzias latipes*.

| | | 0 h | 1 h | 3 h | 6 h | 12 h | 24 h |
|---|---|---|---|---|---|---|---|
| Alive | pH | 7.70 | 7.64 | 7.60 | 7.58 | 7.78 | 7.58 |
| | DO (mg/L) | 7.58 | 7.33 | 6.52 | 5.10 | 5.47 | 5.22 |
| | Temperature (℃) | 24.8 | 24.2 | 24.1 | 24.1 | 24.0 | 23.9 |
| Dead | pH | 7.27 | 7.57 | 7.08 | 7.59 | 7.53 | 7.16 |
| | DO (mg/L) | 7.24 | 6.59 | 6.52 | 6.43 | 5.71 | 0.45 |
| | Temperature (℃) | 25.1 | 23.9 | 23.9 | 23.9 | 23.9 | 23.9 |

**Supplementary Table S3.** Water quality parameters over time in the eNA decay experiment.

| | | **0 h** | **3 h** | **6 h** | **12 h** | **24 h** | **36 h** | **48 h** | **72 h** | **96 h** |
|---|---|---|---|---|---|---|---|---|---|---|
| 4 ºC | pH | 7.08 | 7.22 | 7.21 | 7.34 | 6.44 | 7.12 | 7.05 | 7.04 | 7.61 |
| | DO (mg/L) | 6.01 | 7.24 | 8.14 | 8.63 | 8.81 | 9.19 | 8.13 | 6.08 | 8.20 |
| | Temperature (ºC) | 24.1 | 5.9 | 5.9 | 3.8 | 4.2 | 4.1 | 3.8 | 3.6 | 3.4 |
| 14 ºC | pH | 7.47 | 7.45 | 7.62 | 7.85 | 6.14 | 6.86 | 6.21 | 6.07 | 5.97 |
| | DO (mg/L) | 6.89 | 7.18 | 7.73 | 7.50 | 7.62 | 7.65 | 6.61 | 6.41 | 7.81 |
| | Temperature (ºC) | 22.9 | 15.2 | 14.8 | 14.1 | 13.8 | 13.8 | 13.8 | 13.8 | 14.1 |
| 24 ºC | pH | 7.05 | 7.03 | 7.03 | 7.12 | 6.30 | 6.90 | 7.02 | 7.43 | 7.42 |
| | DO (mg/L) | 6.53 | 6.78 | 6.82 | 6.94 | 6.82 | 6.83 | 6.66 | 7.00 | 6.58 |
| | Temperature (ºC) | 24.0 | 24.1 | 24.0 | 24.0 | 24.0 | 24.0 | 24.0 | 24.0 | 24.0 |
| 34 ºC | pH | 7.31 | 7.47 | 7.65 | 7.82 | 7.04 | 7.57 | 7.07 | 6.63 | 7.39 |
| | DO (mg/L) | 6.58 | 6.48 | 5.22 | 4.90 | 5.95 | 6.60 | 5.92 | 6.75 | 6.35 |
| | Temperature (ºC) | 24.0 | 34.0 | 34.1 | 34.0 | 34.0 | 34.0 | 34.0 | 34.0 | 32.8 |

At time 0 ($t = 0$), water temperature was measured immediately after the aquaria were transferred to the temperature-controlled water baths.
Thereafter, the water temperature gradually transitioned to the target setpoint temperature.

**Supplementary Table S4.** Summary of post-hoc Holm test results from a linear mixed model comparing decay rate constants between eRNA and eDNA across temperatures.

| Temperature (°C) | Contrast | Estimate | SE | df | t-ratio | *P* value |
|---|---|---|---|---|---|---|
| 4 | eDNA - eRNA | 0.00345 | 0.00519 | 168 | 0.665 | 0.5072 |
| 14 | eDNA - eRNA | 0.01323 | 0.00519 | 168 | 2.551 | 0.0116 |
| 24 | eDNA - eRNA | 0.02879 | 0.00519 | 168 | 5.551 | <0.0001 |
| 34 | eDNA - eRNA | 0.04967 | 0.00519 | 168 | 9.576 | <0.0001 |
